# FLIP is essential for oncogenic *KRAS-*driven lung cancer

**DOI:** 10.64898/2026.08.04.738686

**Authors:** Claudia Hamilton, Shaun Sharkey, Hajrah Khawaja, Matilda Downs, Christopher McLaughlin, Connor N Brown, Declan Doherty, Jennifer Fox, Mihaela Pettigrew, Karl Butterworth, Aaron Phillips, Donna Small, Timothy Harrison, Catherine Higgins, Emma M Kerr, Daniel B Longley

**Affiliations:** Johnston Cancer Research Centre, Queen’s University Belfast, Northern Ireland, UK

## Abstract

Mutations in *KRAS* represent the most common oncogenic event in human cancer and occur in approximately 30% of lung adenocarcinomas. The mechanisms by which lung tumours evade apoptosis induced by oncogenic KRAS-driven stress remain incompletely understood. Here, we identify the anti-apoptotic regulator FLIP (*CFLAR*) as a critical dependency in KRAS-mutant lung cancers. We demonstrate that KRAS-mutant human lung cancer cell lines exhibit elevated FLIP expression and enhanced dependence on FLIP for survival compared to KRAS wild-type counterparts. Subsequently, using genetically engineered mouse models (GEMMs), we show that FLIP is essential for *Kras*-driven lung tumour development *in vivo*. *In vitro,* FLIP-deficient lung cancer cells display spontaneous, caspase-8- dependent apoptosis and hyper-sensitivity to the immune/inflammatory cytokines TNFα and TRAIL. Strikingly, FLIP-null lung cancer cells fail to engraft even in highly immunodeficient orthotopic models that lack TRAIL-expressing immune cells but retain TNFα-expressing monocytes. Moreover, silencing of TNFR1 or TNFα but not TRAIL-R2 rescued constitutive caspase-8-dependent apoptosis in FLIP null lung cancer cells, implicating TNFα/TNFR1 in mediating this apoptotic response. Mechanistically, we find that mutant KRAS sustains FLIP expression via ERK1/2 signalling, thereby protecting cells from caspase-8 activation. Notably, KRAS inhibition downregulates FLIP, sensitising cells to TNFα- and TRAIL-induced apoptosis. These findings uncover a novel KRAS–ERK–FLIP axis that protects tumour cells from caspase-8-mediated apoptosis and reveal FLIP as a key survival factor co-opted by *KRAS*-mutant lung cancers. Beyond identifying FLIP as a promising therapeutic target in *KRAS* mutant lung cancer, our work also provides mechanistic insight into the pro-apoptotic effects of KRAS inhibitors and suggests that FLIP expression may serve as a predictive biomarker to enhance patient stratification and the therapeutic efficacy of these agents in lung cancer.

## INTRODUCTION

*Kirsten rat sarcoma viral oncogene homologue* (*KRAS*) encodes a small GTPase protein that regulates a number of downstream signalling networks including MAPK and PI3K/AKT. Activation of KRAS is regulated by the guanosine triphosphate (GTP) hydrolysis cycle in which, aided by small GTPase-activating proteins (GAPs), KRAS GTPase activity converts GTP to GDP to terminate KRAS signalling(1). *KRAS* is the most commonly mutated oncogene in human cancer, including ∼30% of lung adenocarcinoma cases(2, 3). Almost all oncogenic *KRAS* mutations affect exon 2 codon 12 or 13 and play an active role in tumourigenic processes by either impairing KRAS GTPase activity or disrupting the interaction of KRAS with GAPs. In both cases, the abundance of ‘active’ GTP-bound KRAS is increased, resulting in constitutive activation of downstream pathways(1).

Multiple lines of evidence support oncogenic KRAS signalling as an initiation event in lung cancer. *KRAS* mutations are frequent in lung hyperplasia, the precursor lesion to lung cancer(4–8), demonstrating it is an early event in lung tumourigenesis. Expression of mutant *KRAS* in normal cells is sufficient to drive oncogenic transformation(7, 9). However, studies in genetically-engineered mouse models (GEMMs) highlight that the progression to advanced cancer requires cooperating events, including loss/inactivation of tumour suppressors such as TP53(10), STK11/LKB1(11), SMAD4(12) or CDKN2A(13), that enable escape from oncogenic KRAS-driven stress and thereby permit full malignant transformation. Moreover, in advanced lung cancers, KRAS signalling continues to be essential as demonstrated by shRNA-mediated KRAS depletion(14) or KRAS transgene inactivation(15).

The mechanisms by which cancer cells survive persistent oncogenic KRAS induced stress is not yet fully understood. Cellular FLICE (Fas-associated death domain (FADD)-like IL-1β-converting enzyme)-like inhibitory protein (FLIP) is a non-redundant regulator of caspase-8 that regulates extrinsic cell death activation induced by the TNF, FAS, and TRAIL receptors when these receptors are bound by their ligands, which are typically expressed by immune/inflammatory effector cells(16). FLIP is expressed as short (FLIP(S); called FLIP(R) in mice) and long (FLIP(L))(17, 18) splice forms. By forming inactive heterodimers with procaspase- 8, FLIP(S)/(R) directly inhibit caspase-8 activation at the death-inducing signalling complexes (DISCs) formed by activated death receptors(19). FLIP(L) also forms heterodimers with procaspase-8 at the DISC; however, these heterodimers are catalytically active and whether they inhibit or amplify the apoptotic signal is dependent on their relative abundance, with the high FLIP(L) levels that are typically found in many cancers blocking caspase-8 activation(20, 21). In addition to apoptosis, FLIP has been linked to regulation of autophagy (via binding to ATG3 and impeding its interaction with LC3(22)) and necroptosis (the FLIP(L)/caspase- 8 heterodimer can cleave the necroptosis effectors RIPK1 and RIPK3(23, 24)).

FLIP is a key regulator of cell fate that facilitates cell survival in response to a plethora of cellular stresses, including chemotherapy (25–28), radiotherapy(29), targeted agents(30–35) and immunotherapies(36), and we and others have shown that FLIP is frequently overexpressed in non-small cell lung cancer (NSCLC) and is predictive of poorer treatment and survival outcomes(34, 37, 38). Here, we define the role of FLIP in the development, progression and maintenance of *KRAS*- driven lung adenocarcinoma.

## MATERIALS AND METHODS

### Compounds

Recombinant Mouse TRAIL (TNFSF10 Protein) TNFα were purchased from BioTechne, Emricasan, AZD5991 and LY3214996 from Selleckchem (Waltham Abbey, UK), MRTX1133 from MedChemExpress (NJ, USA) and PF-4708671 from Sigma (Dorset, UK). Cisplatin was obtained from Belfast City Hospital Pharmacy.

### Mice studies, adenoviral infection and *in vivo* Cre-recombination

Cflar Flox (Strain #:022009, allele: Cflar^tm1Ywh^/J (39) and p53 Flox (Strain #:008462, allele: Trp53tm1Brn/J (40)) mice were purchased from Jax. Kras LSLG12D (Jax Strain #:008179, allele: Krastm4Tyj/J, (6)) were provided by Sansom Lab (Scotland Institute). Kras LSLG12D and p53 Flox mice were bred to generate KP mice (Kras LSLG12D/WT;p53Flox/Flox), which were then crossed with Cflar Flox mice to generate KP (Cflar +/+) and KPF (Cflar Fx/Fx) founding pairs and bred for experiment. All mice were genotyped by Transnetyx (Cordoba, TN, USA). All experimental procedures were performed in accordance with the Department of Health NI regulations (Project Licences 2810 and 2874) and were approved by the Queen’s University Belfast’s Animal Welfare and Ethical Review Committee (AWERB). Endogenous lung tumours were induced in *Kras^G12D/+^;Trp53^Fx/Fx^ and Kras^G12D/+^;Trp53^Fx/Fx^;Cflar^Fx/Fx^* mice between 8-12 weeks of age by intranasal delivery of adenoviral Cre-recombinase (5x10^7^ PFU per mouse, (University of Iowa Vector Core)), as previously described(41). For transplantation studies in both immunocompetent (C57Bl/6J; Inotiv, UK) and immunocompromised mice (C57BL/6N-*Rag2^Tm1^-IL2rg^Tm1^*/Rj; Janvier, France), 5x10^5^ cells in PBS were implanted via tail vein injection. Mice were monitored for signs of bodyweight loss and clinical symptoms (piloerection, hunching, laboured breathing, and reduced movement).

Animals were culled and considered to have met the scientific endpoint when presenting with clinical symptoms and bodyweight loss simultaneously. Lungs were inflated using PBS and fixed in 10% formalin overnight at room temperature, prior to dehydration and embedding.

### CBCT scans

Preclinical Cone Beam Computed Tomography (CBCT) scans were acquired on the Small Animal Radiation Research Platform (SARRP, Xstrahl Life Sciences, UK) at 10-, 12- and 14-weeks post tumour initiation, using an imaging energy of 60 kV with a 0.5 mm Al filter. Image reconstruction was performed using 360° images and filtered back-projection without postfiltering. Log(white/x) was applied to input images using FDK with a Hamming filtering window. CBCT scans had a slice thickness of 0.26mm and a voxel size of 0.275 mm. Manual contours of the lungs were created by a blinded observer using Muriplan© (Xstrahl Life Sciences). Manual contours were created slice-by-slice in the coronal plane, with corrections made using other planes. Tumour volume was calculated from the manual contours using the dose volume histogram calculation in Muriplan©.

### Histology and tumour grading

Histological analysis was carried out on formalin-fixed, paraffin-embedded lung tissue sections. Tumour burden was assessed from haematoxylin and eosin (H&E) stained sections using QuPath® software. The total tumour tissue was measured (µm^2^) and calculated as a percentage of total lung area on the section. Tumour grade, categorised as hyperplasia, adenoma or adenocarcinoma, was assessed on H&E-stained sections as previously described(6).

### Generation of cell lines

Lung tumour cell lines were generated from tumour-bearing *Kras^G12D/+^;Trp53^Fx/Fx^;Cflar^Fx/Fx^* mice. Tumours were collected in ice-cold Hank’s balanced salt solution (HBSS), dissected into smaller pieces and digested using Collagenase/Dispase (Roche, Mannheim, Germany) for 2.5 h at 37°C. HBSS containing 10% FBS and DNAse (1 mg/30 mL) was added, large tissue was removed through 70 µm filtration and cells were plated in complete media (DMEM/F12 containing 10% FBS, 4 mM _L_-glutamine, and 1% penicillin/streptomycin) for 90 minutes. The media (containing tumour-enriched slower adhering cells) was then replated. Cells were cultured in DMEM/F12 media (Gibco, Loughborough, UK) supplemented with 10% FCS and 1% penicillin/streptomycin and passaged every 3-5 days. All cells were maintained at 37 °C in a 5% CO2 humidified atmosphere and were regularly screened for the presence of mycoplasma using the MycoAlert Mycoplasma Detection Kit (Lonza, Slough, UK).

### Genotyping

DNA was extracted from organoids and cell lines using DNeasy Blood and Tissue Kit (Qiagen 69504, Manchester, UK) according to manufacturer’s instructions. Recombination of Kras and Trp53 was confirmed using the primers and protocols described (https://jacks-lab.mit.edu/protocols.html). To determine the presence of Cflar floxed or WT alleles 250ng of genomic DNA was amplified by PCR for 30 cycles at 94 °C for 30s, at 58 °C for 60s, and 72 °C for 120s. PCR amplicons were separated on a 2% (w/v) agarose gel (336bp Cflar WT, 500bp Cflar^Fx^(39)).

### *In vitro* Cre-recombination

*In vitro* recombination was carried out using adenovirus expressing Cre recombinase and EGFP (Ad5CMVCre-eGFP (approximate titer 1x10^10^-7x10^10^ PFU/mL) (University of Iowa, Viral Vector Core)) (AdV-Cre). Control cells were infected with Ad5CMVempty (University of Iowa, Viral Vector Core) (AdV-EV). For generation of stable FLIP KO lines (KPF3/KPF4), cells were seeded at a low seeding density in p90 plates. AdV-EV and AdV-Cre viral suspensions were prepared in EMEM (Sigma Aldrich). Viral media was added to cells at a final dilution of 1:1000 virus:media. Clones were isolated using Pyrex cloning cylinders (Corning) and grown to confluency before transferring to culture flasks. Knockout was confirmed by western blotting and PCR.

### siRNA transfection

Pre-designed siRNAs were purchased for Caspase-8, Tnfrsf10b (DR5), Tnfα and Tnfrsf1a (TNFR1) (SMARTpool, Dharmacon, Cambridge, UK). Transfection of KPF cells was carried out using Lipofectamine RNAiMAX (Invitrogen) according to the manufacturer’s instructions.

### Western blotting

Whole cell lysate from cell lines and organoids was prepared in RIPA buffer containing protease inhibitor (539134-1SET, Merck) and phosphatase inhibitor (Roche) and quantified by BCA assay. The following primary antibodies were purchased from Cell Signalling Technology: FLIP (D5J1E) #56343), cleaved caspase-8 (Asp374; #9496), Caspase-3 (#9662S), pAKT^S473^ (#4060), AKT (#9272), phospho-p44/42 MAPK (ERK1/2^T202/Y204^) (#9101), p44/42 MAPK (ERK1/2) (#9202), phospho-S6^S235/6^ ribosomal protein (91B2) (#4857), S6 ribosomal protein (54D2) (#2317), TNFα (#3707). DR5 antibody (SC-57086) was purchased from Santa Cruz (TX, USA), murine KRAS clone 2C1 antibody (LS- C175665-LSP) from LifeSpan Biosciences (CA, USA) and β-actin (A5316) antibody from Sigma. The secondary antibodies used for Western blot detection were goat anti-rabbit and anti-mouse IgG- HRP conjugates (Cell Signalling Technologies, MA, USA)). Protein expression was detected using the Western Lighting Plus ECL (PerkinElmer, CT, USA) or WesternBright Sirius HRP substrate (Advansta, CA, USA) on the G:BOX Chemi6 gel doc system (Syngene, Cambridge, UK).

### Caspase activity assay

Caspase activity was assessed using Caspase-3/-7 Glo Luminescent Assay (Promega, Southampton, UK) according to manufacturer’s instructions.

### qPCR

RNA was extracted using High Pure RNA Isolation kit (Roche) according to the manufacturer’s instructions. cDNA was synthesized using Transcriptor First Strand cDNA synthesis kit (Roche). qPCR was carried out using SYBR green (Roche) on LC480 light cycler (Roche) according to the manufacturer’s instructions.

### High content microscopy

Cells were seeded onto black, glass-bottomed 96-well plates (Cellvis, CA, USA) at appropriate densities and treated as required. At the experimental endpoint cells were stained with FITC-tagged Annexin-V (BD-Biosciences), 0.333 µg/mL propidium iodide (Sigma Aldrich), and 1.33 µg/mL Hoechst (Invitrogen). High content screening was performed using the Array Scan XTI high-content microscope (Thermo Scientific) and analysed by HCS Studio Cell Analysis Software (Thermo Scientific).

### Flow Cytometry

To assess cell death in cells infected with AdV-Cre, live cell staining with APC- tagged Annexin V (BD Biosciences, Berkshire, UK) and 7-Amino-Actinomycin D (7-AAD) (BD Biosciences) were analysed on a BD Accuri C6 Plus flow cytometer (BD Biosciences). For all other cell lines FITC-Annexin V (BD Biosciences) and Propidium iodide (Merck) staining was used.

### Cell Viability Assay

Cell viability was assessed using CellTiterGlo Luminescent Assay (Promega, Madison, WI) according to manufacturer’s instructions. Three-dimensional (3D) cell viability was determined with CellTiter-Glo 3D Cell Viability Assay (Promega), according to the manufacturer’s instructions.

### Clonogenic Assay

Cells were seeded at 150 cells/well, media and drug was refreshed every 96h. Colonies were fixed with MeOH and stained with crystal violet after 8 days treatment.

### Statistical analyses

Statistical analyses were performed using Prism 10.0 software (Graphpad). *P*<0.05 was considered significant. Results between two groups were compared using two-tailed Student’s *t*-test. One-way ANOVA was used for analyses of three or more groups and adjusted for multiple comparisons with Dunnetts correction. Ordinary two-way ANOVA was used for analyses that involved two variables with Bonferroni post-hoc tests for multiple comparisons. Kaplan-Meier comparison was used for analysis of survival cohorts and Fisher’s exact test was used to analyse genotyping data.

## RESULTS

### *KRAS* mutation correlates with high FLIP expression and dependence in human lung cancer cells

To investigate the relationship between oncogenic KRAS signalling and FLIP expression in human cancer, we assessed FLIP dependence in the Broad Institute DepMap database (https://depmap.org/portal/) and found that *KRAS* mutant (MT) models are significantly more FLIP-dependent than *KRAS* wild-type (WT) cell lines across all solid cancer cell lines (p < 0.0001; **Figure 1A**) and specifically in lung cancers (p < 0.01; **Figure 1B**). Moreover, we found that FLIP expression is significantly elevated in *KRAS* MT lung cancer cell lines versus *KRAS* WT counterparts (**Figure 1C**); indeed, this was observed across all solid tumour cell lines (**Supplementary Figure 1A**). Notably, we found a strong correlation between FLIP-dependence and FLIP mRNA expression (Spearman r = -0.47, p < 0.0001; **Figure 1D**), indicating that FLIP dependence is higher in cell lines with high FLIP expression; again, this correlation held up across all solid tumour cell lines (Spearman r = -0.51, p < 0.0001; **Supplementary Figure 1B**). Expression of FLIP’s binding partner and paralog procaspase-8 was also found to be significantly higher in *KRAS* MT cell lines than *KRAS* WT lung cancer cell lines (p < 0.0001; **Figure 1E**), and, consistent with FLIP’s canonical mechanism-of-action, a significant correlation was also observed between FLIP-dependence and expression of procaspase-8 (Spearman r = - 0.62, p < 0.0001; **Figure 1F**); similar correlations were noted across all solid tumour lines (**Supplementary Figure 1C/D**). These data reveal that *KRAS* MT cancer cells are more dependent upon FLIP for survival, and FLIP or procaspase-8 expression can be used as a surrogate indicator of this dependency.

**Fig. 1:**
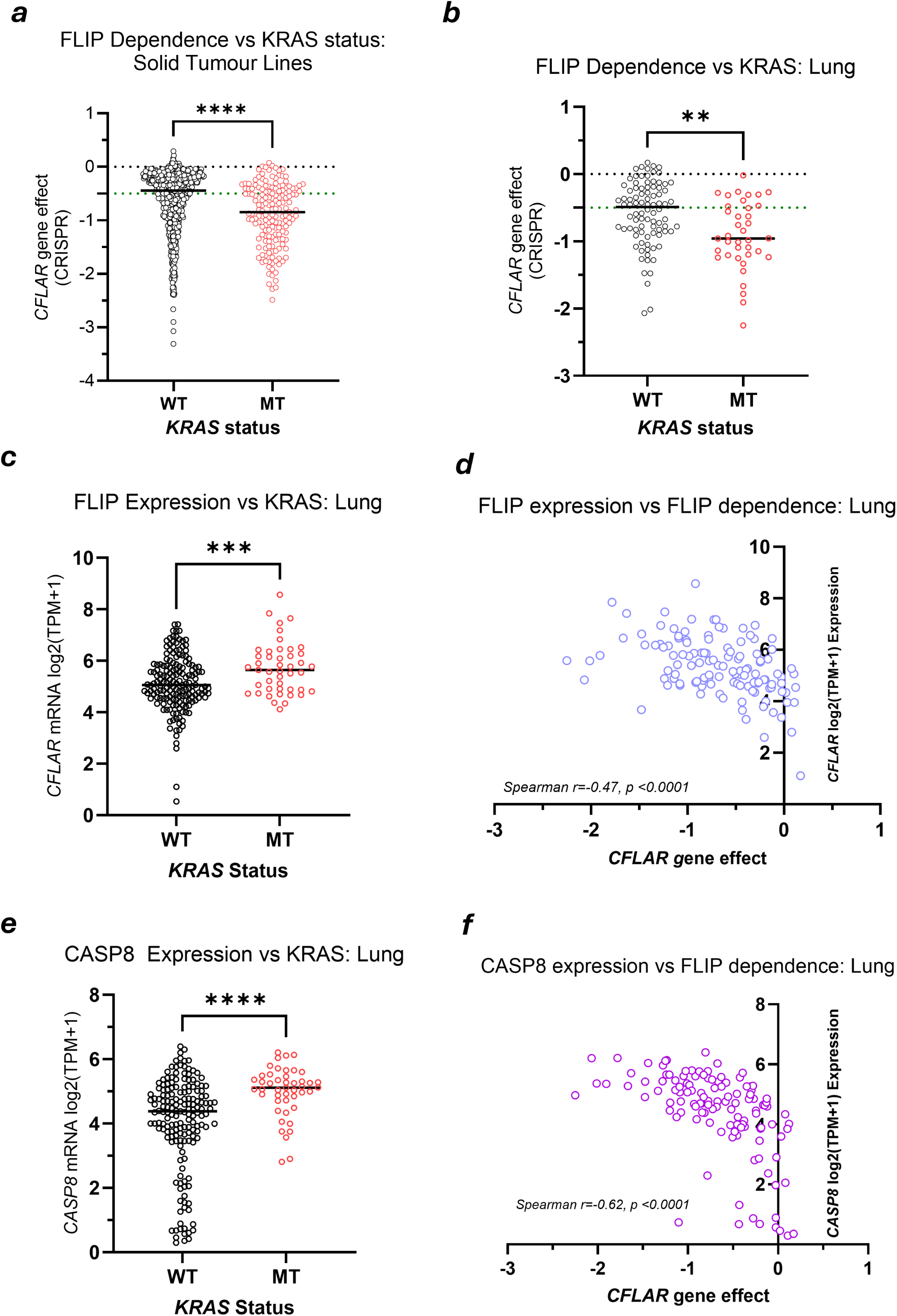
*KRAS* mutation correlates with high FLIP expression and dependence in lung cancer cells. (**A**) Dependence of *KRAS* WT and MT solid cancer cell lines on FLIP/*CFLAR* assessed using the CRISPR DepMap data (Public 24Q4+Score, Chronos). (**B**) Dependence of *KRAS* WT and MT lung cancer cell lines on FLIP/*CFLAR* assessed using the CRISPR DepMap data (Public 24Q4+Score, Chronos). (**C**) Expression of FLIP/*CFLAR* mRNA in *KRAS* WT and MT lung cancer cell lines in the DepMap lung cancer cell line panel. (**D**) Correlation of FLIP/*CFLAR* gene expression with FLIP/*CFLAR* gene dependency in lung cancer cell lines in the DepMap panel. (**E**) Expression of Caspase-8/*CASP8* mRNA in *KRAS* WT and MT solid cancer cell lines in the DepMap lung cancer cell line panel. (**F**) Correlation of Caspase-8/*CASP8* gene expression with FLIP/*CFLAR* gene dependency in lung cancer cell lines in the DepMap panel.

### FLIP-deficient murine *KRAS* mutant/p53 Null lung cancers fail to grow *in vivo*

Collectively, the results presented in **Figure 1** suggest that FLIP is required to maintain the viability of *established KRAS* MT lung cancer cells. As *KRAS* mutations are an early driving event in lung cancer transformation, we investigated the dependence on FLIP of oncogenic KRAS-driven lung tumour initiation. To do this, we crossed the established *Kras* G12D/*Trp53* null (KP) mouse model(10) with mice in which the FLIP gene (*Cflar*) is “floxed” around exon 1(39) (**Fig. 2A**), to generate *Kras^G12D/+^;Trp53^Fx/Fx;^Cflar^Fx/Fx^* (KPF) mice. Lung tumours were induced by intranasal delivery of adenoviral Cre (AdV-Cre) recombinase in KP and KPF mice, and longitudinal Computed Tomography (CT) analysis of tumour burden over 10- 14 weeks demonstrated significant increases in lesion volume over time in KP mice as expected, but no significant change in volume in KPF mice, indicating no detectable or very low tumour burden (**Fig. 2B/C**). In light of these results, we performed a survival study. In keeping with the well characterised timeline for KP mice, results showed that KP animals reached clinical endpoint between 14-16 weeks post AdV-Cre; however, 100% of KPF mice were still alive 6 weeks later (153 days post-delivery of AdV-Cre), at which point animals were culled and the difference in survival analysed by Kaplan-Meier log-rank test (**Fig. 2D**). We also analysed lung tumour burden by H&E staining 16 weeks after tumour initiation, graded as previously described(42): KP mice had high tumour burden (mean = 37 lesions/mouse) with hyperplasias, adenomas and adenocarcinomas all evident; whereas KPF mice had no or very few lesions evident (mean 5 lesions/mouse), and detected disease was limited to hyperplasias (**Fig. 2E**). These results indicate that FLIP is required for establishment and/or growth of mutant Kras-driven lung cancer.

**Fig. 2:**
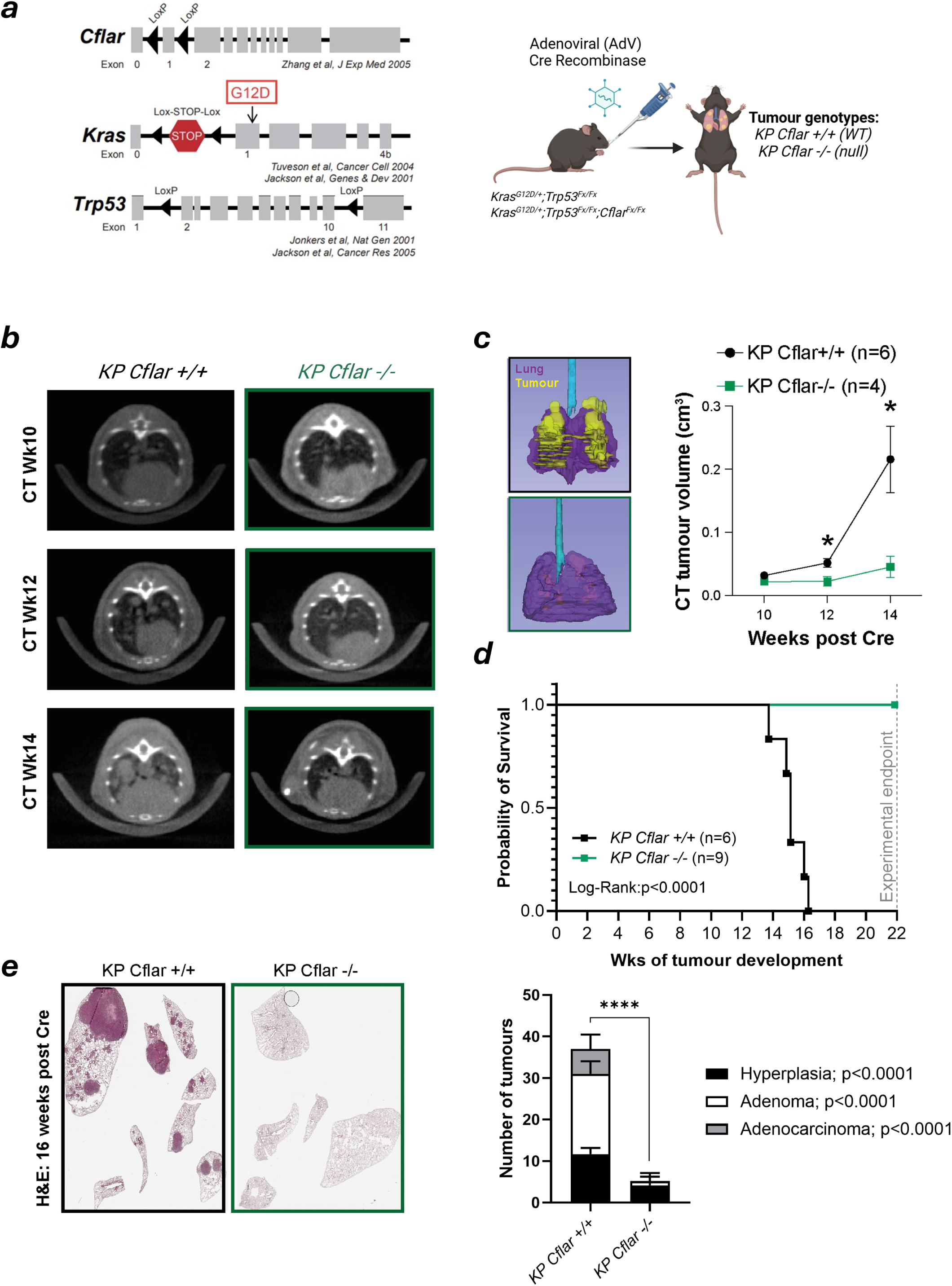
FLIP/*Cflar* is essential for *Kras*-driven lung tumour development. (**A**) Schematic depicting the induction of tumours in transgenic *Kras^G12D/+^;Trp53^Fx/Fx^* (*KP Cflar* +/+ (*WT))* or *Kras^G12D/+^;Trp53^fx/fx^;Cflar^fx/fx^* (*KP Cflar* -/- *(null))* mice by intranasal delivery of Adenoviral (AdV) Cre recombinase. (**B**) Representative CT scans from KP *Cflar +/+* and KP *Cflar -/-* mice at week 10 (Wk10), 12 (Wk12) and 14 (Wk14) post administration of Cre recombinase. (**C**) 3D rendering of lungs from a KP *Cflar +/+* and KP *Cflar -/-* mouse from micro-CT scans. Lung tissue is highlighted in purple; tumours are highlighted in yellow. CT tumour volume (cm^3^) was calculated from manual contours of lung CT scans from (**B**) using Muriplan©. Significance was determined by Students *t* test, *p<0.05. (**D**) Kaplan-Meier curve showing the survival of KP *Cflar* +/+ (n=6) vs KP *Cflar* -/- (n=9) mice. Survival analysis was performed using Mantel-Cox log-rank test. (**E**) Representative H&E and quantification of lung tumours from KP *Cflar +/+* and KP *Cflar -/-* mice at 16 weeks post-Cre. Data are grouped by histologic tumour grades (hyperplasia, adenoma, adenocarcinoma). Significance was determined by Students *t* test, ****p<0.0001. Data are the mean +/- SD.

### FLIP-deficient lung cancer cells have elevated caspase-8 and apoptosis rates

To explore the mechanism behind the failure of FLIP-deficient Kras-driven lung cancers to establish *in vivo* (**Figure 2**), we induced lesions in KPF mice as above and aged animals until they displayed clinical symptoms of lung tumour burden ∼30 weeks post adenoviral Cre delivery. We isolated epithelial tumour cell lines from KPF tumours using differential detachment as previously described(43) (**Fig. 3A**). Assessment of the genotype of these *ex vivo* lung cancer cell line models indicated that although they showed efficient recombination of *Kras^G12D^* and *Trp53* alleles, remarkably, they all still retained FLIP (*Cflar*) (**Fig. 3B/C**), further underlining the essentiality of FLIP in Kras-driven lung cancer, and indicating that FLIP remains essential in established lung tumours (as suggested by the DepMap data in human models, **Figure 1**). To explore the intrinsic dependence of established KP lung cancer cells on FLIP, we transduced the isolated lines with AdV-Cre to induce *Cflar* recombination *ex vivo* and generated clonal cell lines (**Fig. 3D**). Successful recombination was confirmed by PCR (**Fig. 3E**) and Western blot (**Fig. 3F**) in 2 independent clones (“KPF3” and “KPF4”). Notably, compared to cells that retained FLIP (“KPF1”, “KPF2”), knockout of FLIP resulted in increased caspase-8 and caspase-3 activation (**Fig. 3F-H**) and significantly elevated levels of cell death (**Fig. 3I**), confirming FLIP’s role in suppressing apoptosis in this model. In addition, the elevated cell death in the KPF3 and KPF4 models was found to be caspase-8-dependent (**Fig. 3J**), and siRNA-mediated caspase-8 depletion rescued the cell death induced by acute Cre-mediated FLIP-deletion in KPF1 and KPF2 models (**Fig. 3K**). Together, these data illustrate that FLIP remains essential in established *Kras* mutant lung cancers, in which it is required to suppress caspase-8-mediated cell death.

**Fig. 3:**
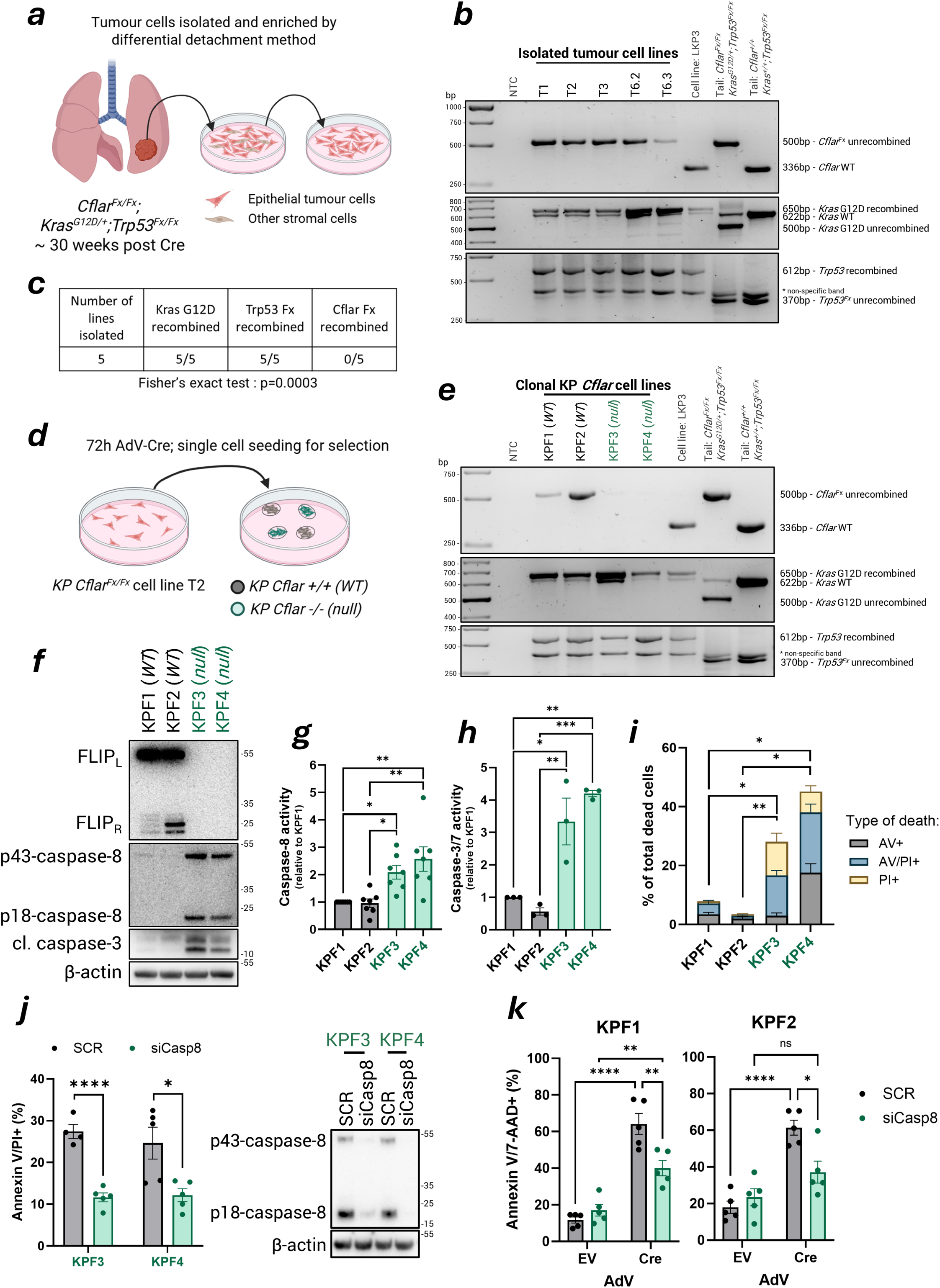
FLIP-deficient lung cancer cells have elevated caspase-8 and apoptosis rates. (**A**) Schematic representing the generation of *Cflar^Fx/Fx^ Kras^G12D/+^ Trp53^Fx/Fx^* cell lines. (**B**) PCR analysis of *Cflar, Kras, Trp53* loci in isolated tumour cells lines showing the presence of the unrecombined Cflar^Fx^ 500bp amplicon and recombination of *Kras* G12D and *Trp53*. Each lane (T1-T6) are cell lines generated from lung tumours of individual mice, lanes T6.2 and T6.3 are cell lines generated from individual tumours in the same mouse. DNA from matched genotype mouse tail (*Cflar*^Fx/Fx^; *Kras* G12D/+; *Trp53*^Fx/Fx^) was used as a negative control. KP3 cells were used as a positive control for *Kras* G12D and *Trp53*^Fx/Fx^ recombination. (**C**) The occurrence of recombination of the *Kras, Trp53* and *Cflar* loci in each cell line was compared using Fishers exact test, p=0.0003. (**D**) Experimental schematic showing the generation of individual KP *Cflar* -/- (*null*) and KP *Cflar +/+* (*WT*) clones by transduction of *KP Cflar*^Fx/Fx^ cell line T2 with adenoviral Cre (AdV-Cre). (**E**) PCR analysis of *Cflar, Kras* and *Trp53* loci in KPF WT and null cell lines showing the absence of the unrecombined *Cfla*r^Fx^ 500bp amplicon in KPF null cells (KPF3/KPF4). (**F**) Western blot analysis of basal expression of FLIP_L_, FLIP_R_, cleaved caspase-8 (p43 and p18) and cleaved (cl.) caspase-3 in KPF cell lines. β- actin was used as a loading control. Basal activity of activator caspase-8 (**G**) and effector caspase-3 and –7 (**H**) was analysed in extracted protein lysates from KFP cell lines (n>/=3 per biological replicates per cell line) using CaspaseGlo® activity assays. Each cell line is presented relative to the activity of KPF1 cells. (**I**) Basal cell death in KPF cell lines assessed at 48 hours post seeding by Annexin V/PI staining. Bar graph shows the percentage dead cell population for each cell line (n=3). (**J**) Annexin V/PI analysis in KPF3 and KPF4 cells transfected with control (SCR) and *Casp8* siRNA for 72 hours (n=4). Results were compared using by Students *t*-test. Matched western blot analysis of p43/p18-caspase-8 and β-actin in KPF3 and KPF4 confirming silencing of caspase-8. (**K**) Annexin V/7-AAD analysis in KPF1 and KPF2 cells following simultaneous transduction with adenoviral Cre recombinase (AdV-Cre) or empty vector (AdV-EV) and transfection with control (SCR) and Casp8 siRNA for 48 hours (n=5). Statistical tests were performed using one-way ANOVA and adjusted for multiple comparisons with Dunnetts correction (G-I) and two-way ANOVA with Bonferroni correction (K). (\**p <* 0.05, \*\**p <*0.01, **** p<0.0001, ns – non-significant). Data are mean +/- SEM.

### FLIP-deficient cells grow and form colonies *in vitro* but fail to establish *in vivo*

Although undergoing significant cell death (**Fig. 3I**), the FLIP deficient KPF3 and KPF4 lung models were able to be cultured and form colonies; indeed surprisingly, the KPF4 model was as proficient as KPF1 and KPF2 models in forming colonies, and both FLIP-null models had similar doubling times as the FLIP-proficient models (**Fig. 4A** and **Supplementary Fig. 2A**) despite their elevated caspase activity and rates of apoptosis. Moreover, the KPF3 and KPF4 FLIP null models maintained elevated caspase-8 and caspase-3/7 activity through multiple passages (**Fig. 4B** and **Supplementary Fig. 2B**), suggesting that although FLIP loss *in vitro* leads to elevated caspase activity and enhanced rates of cell death, sufficient numbers of cells in the overall clonal population are able to survive and proliferate. However, when FLIP KO KPF4 cells were inoculated into the tail vein of immune-proficient mice to seed lung tumours, all of the mice survived until the study endpoint (13 weeks post-inoculation), whereas animals inoculated with FLIP-proficient KPF2 cells reached survival endpoint as expected ∼70-90 days post injection (**Fig. 4C**). Moreover, zero tumours were found in the mice inoculated with FLIP KO KPF4 cells, whereas mice inoculated with FLIP-proficient KP cells had highly significant tumour burdens (**Fig. 4D/E** and **Supplementary Fig. 3**). Combined with the *in vitro* data, this suggested that anti-tumour immunity plays a role in preventing FLIP null cells from establishing and growing in the lung. However, when the same experiment was repeated in highly immune-deficient (C57BL/6N-Rag2^Tm1^- IL2rg^Tm1^/Rj) animals, we observed the same phenotype (**Fig. 4F-H** and **Supplementary Fig. 4A**).

**Fig. 4:**
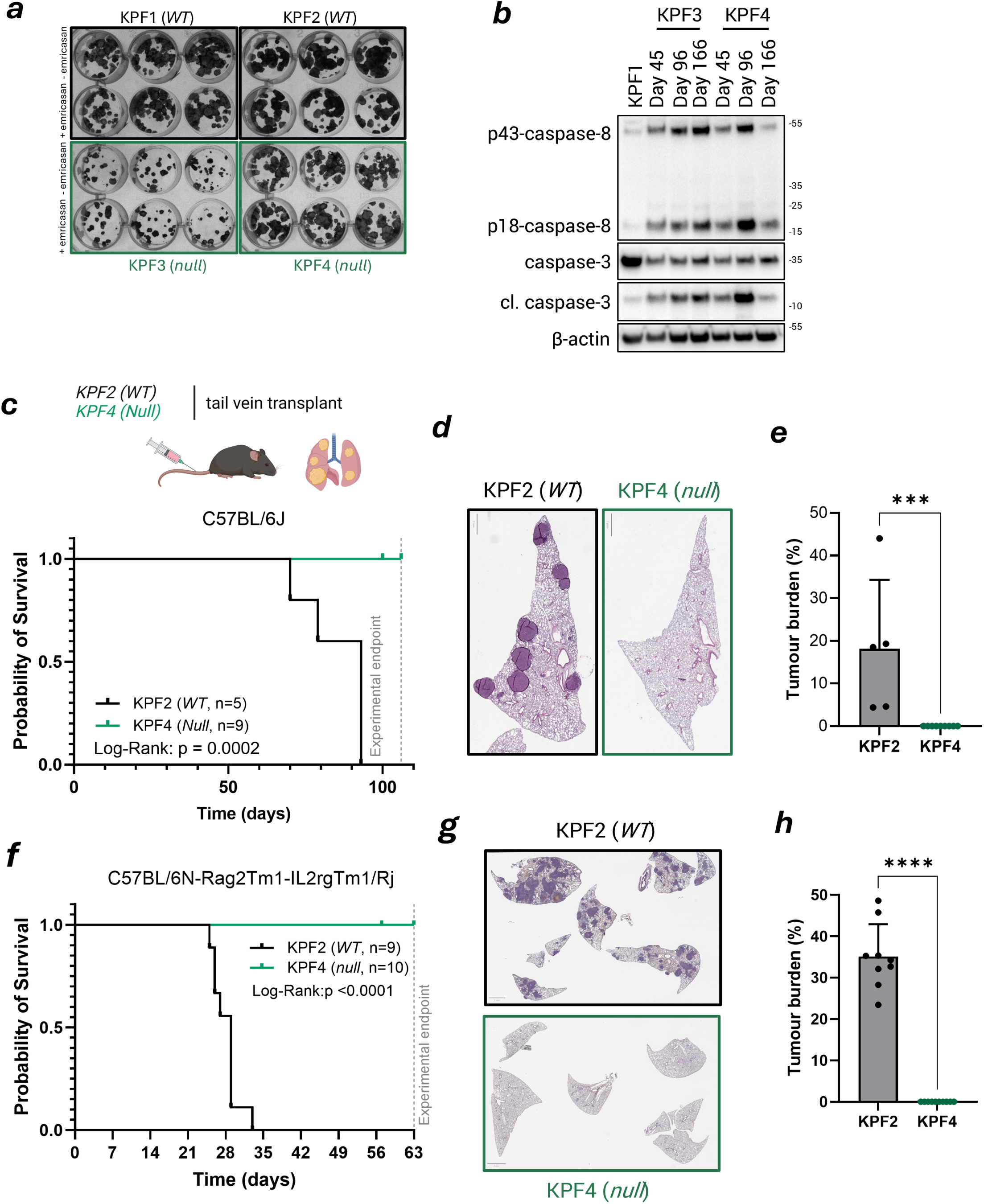
FLIP-deficient cells grow and form colonies *in vitro* but fail to establish *in vivo*. (**A**) Representative images of colony formation in KPF *Cflar WT* (KPF1, KPF2) and *Cflar Null* (KPF3, KPF4) cells in the presence or absence of emricasan (10µM). (**B**) Western blot analysis of basal expression of p43/p18- caspase-8, caspase-3 and β-actin in KPF3 and KPF4 cells at day 45, 96 and 166 after the removal of AdV-Cre. KPF1 was used as a control. (**C**) Experimental scheme showing tail vain transplantation of KPF cell lines in C57Bl/6J mice. Kaplan-Meier curve showing the survival of mice transplanted with KPF2 (*WT*) (n=5) or KPF4 (*null*) (n=9). Survival analysis was performed using Mantel-Cox log- rank test. (**D**) Representative H&E of lung sections from KPF2 (*WT*) and KPF4 (*null*) mice. (**E**) Quantification of histologic tumour burden (tumour area/total lung area) from mice at 13 weeks post-inoculation with KPF2 (*WT*) or KPF4 (*null*) cells. Statistical significance was assessed using the Mann–Whitney test, ****p<0.0001. (**F**) Kaplan-Meier curve showing the survival of C57BL/6N-Rag2^Tm1^-IL2rg^Tm1^/Rj mice following tail vein transplant of KPF2 (*WT*) (n=9) or KPF4 (*null*) (n=10) cells. Survival analysis was performed using Mantel-Cox log-rank test. (**G**) Representative H&E of lung sections from KPF2 (*WT*) and KPF4 (*null*) mice. (**H**) Quantification of histologic tumour burden (tumour area/total lung area) from KP *Cflar +/+* and KP *Cflar -/-* mice at experimental endpoint. Statistical significance was assessed using the Mann–Whitney test, ****p<0.0001. Data are mean +/- SD.

### FLIP-deficient cells are hypersensitive to TRAIL and TNFα but not BH3 mimetics

FLIP is a non-redundant inhibitor of the extrinsic apoptotic pathway, which via BID cross-talks with the intrinsic apoptotic pathway regulated by BCL-2 family proteins (**Fig. 5A**). As expected therefore, we found that FLIP KO models were hyper- sensitive to exogenous recombinant murine TRAIL (**Fig. 5B/C**), whereas the FLIP- proficient cells were highly TRAIL resistant. However, FLIP-null KP cells were *not* more sensitive than FLIP-proficient counterparts to direct activators of the mitochondrial apoptotic pathway (**Figure 5A/D and Supplementary Fig. 2C**). Notably, FLIP-null lung cancer cells *were* more sensitive than control cells to cisplatin (**Fig. 5E**), suggesting that chemotherapeutic DNA damaging agents induce FLIP-dependent apoptosis, consistent with our previous studies (37, 38).

**Fig. 5:**
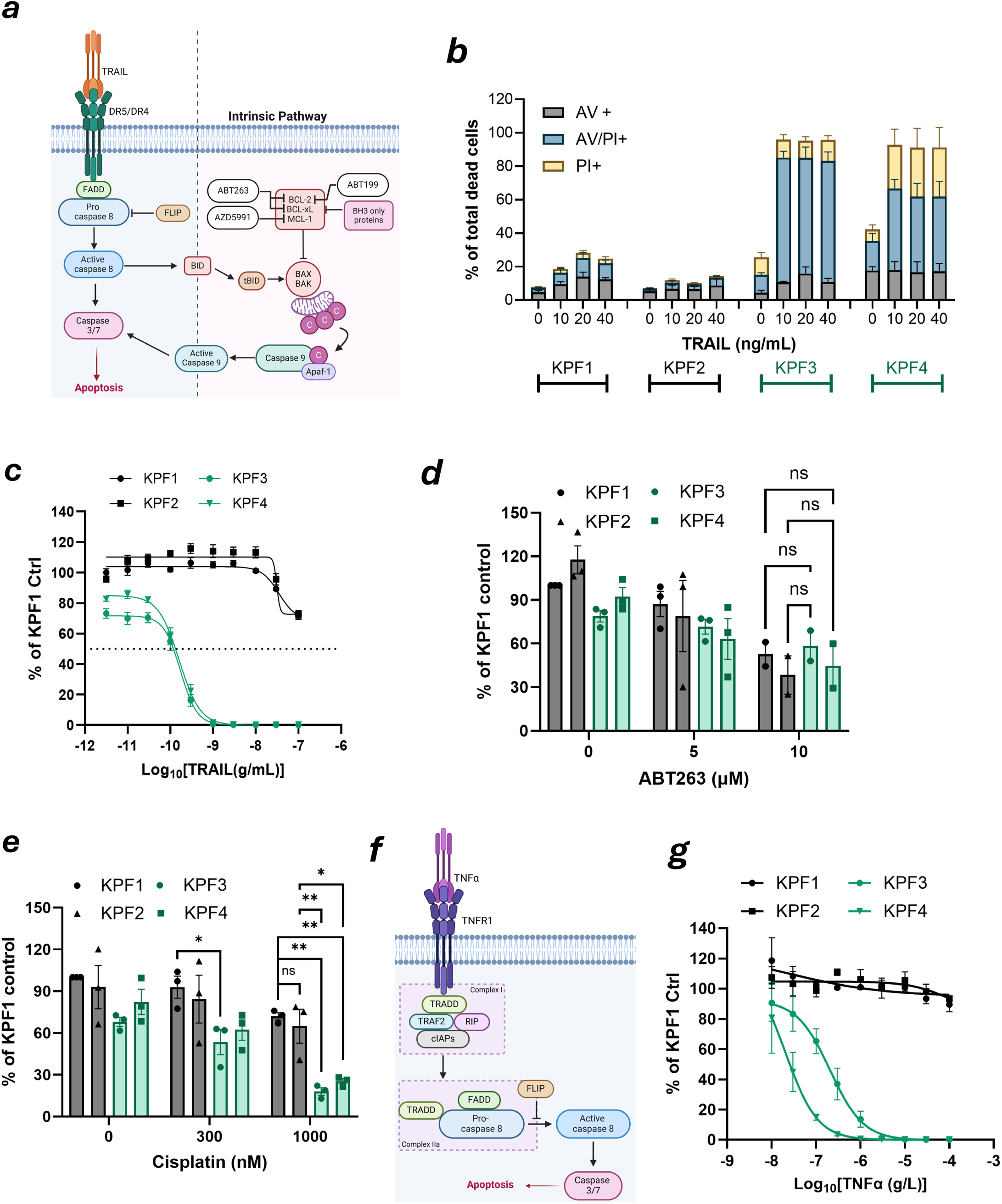
FLIP-deficient cells are hypersensitive to TRAIL and TNFα but not BH3 mimetics. (**A**) Schematic illustrating the link between the death receptor pathway and the intrinsic apoptosis pathway. TRAIL binding to DR5/4 activates a downstream signalling cascade through FADD and procaspase-8. FLIP antagonizes procaspase-8 activation. Active caspase-8 activates both caspase-3/7 and BID. Truncated BID (tBID) translocates to the mitochondria to induce BAX/BAK mediated mitochondrial outer membrane permeabilisation (MOMP). Subsequent cytochrome *c* release prompts assembly of the apoptosome, activation of procaspase-9 and cleavage of caspase-3/7. The BH3 mimetics ABT263, ABT199, AZD5991 inhibit the function of antiapoptotic proteins BCL-2 (ABT199, ABT263), BCL-xL (ABT263) and MCL-1 (AZD5991) to promote BAX/BAK activation. (**B**) Annexin V/PI analysis in KPF1-4 cells following treatment with 10, 20 or 40ng/mL of murine TRAIL for 24 hours, (n=3). (**C**) Cell viability assays in KPF1-4 cell lines following treatment with increasing half-log doses of murine TRAIL (0–100ng/mL). Data was normalized to KPF1 control, (n=3). (**D**) Cell viability assays in KPF1-4 cell lines following treatment with increasing doses (1.25, 2.5, 5 and 10µM) of BH3 mimetics ABT263. Data was normalized to KPF1 control, (n>=2). (**E**) Cell viability assay in KPF1-4 cell lines following treatment with increasing doses (1.25, 2.5, 5 and 10µM) of cisplatin. Data was normalized to KPF1 control. (n=3). Statistical tests were performed using two-way ANOVA and adjusted for multiple comparisons with Bonferroni correction. (*p<0.05\*\**p <*0.01, ns – non-significant). (**F**) Schematic depicting the TNFα death receptor pathway. TNFα ligand binding to TNFR1 activates a downstream signalling cascade through TRADD, TRAF2, RIP and cIAPs (Complex I). Complex IIa assembly follows detachment of TRADD from TNFR1, recruitment of FADD and procaspase-8. Apoptosis follows activation of caspase-3/7 by caspase-8. FLIP antagonizes procaspase-8 activation. (**G**) Cell viability assays in KPF1-4 cell lines following treatment with increasing half-log doses of TNFα (0–100ng/mL). Data were normalized to KPF1 control, (n=3) and are plotted as mean +/- SEM.

TRAIL is predominantly expressed by immune effector cells such as CD8+ T-cells and natural killer cells(44)); however, neither of these cells is present in C57BL/6N- Rag2^Tm1^-IL2rg^Tm1^/Rj animals. This suggested that the *in vivo* FLIP-dependency on FLIP in Kras-driven lung cancers (**Figures 2** and **4**) was not dependent on canonical signalling via TRAIL. TNFα is another immune cytokine capable of activating caspase-8 in a FLIP-dependent manner (**Fig. 5F**), and C57BL/6N- Rag2^Tm1^-IL2rg^Tm1^/Rj animals retain TNFα-expressing monocytic cells(45). Notably, both FLIP KO models were exquisitely sensitive to TNFα *in vitro*, whereas the FLIP- proficient models were completely resistant (**Fig. 5G**).

### FLIP-deficiency drives TNFα /TNFR1-mediated apoptosis in Kras lung cancer

The above results suggested that TNFα/TNFR1 signalling may be responsible for *in vitro* apoptosis and *in vivo* rejection of FLIP null lung cancer cells. In support of this, when we examined the roles of the TRAIL receptor DR5 (mice have only have one TRAIL receptor) and the TNFα receptor, TNFR1, *in vitro* we found that only silencing of TNFR1 reduced apoptosis induction in the KPF1 model in which FLIP was acutely deleted (**Fig. 6A**) and in the KPF4 model that lacks FLIP (**Fig. 6B** and **Supplementary Fig. 4B**). Of note, TNFR1 gene expression was the same across all KPF models (**Supplementary Fig. 4C**), such that differences in TNFR1 expression are not responsible for our *in vivo* observations. Consistent with the importance of TNFR1 for the observed phenotypes, silencing of TNFα also rescued cell death induction and caspase activation in FLIP-deleted KPF1 cells (**Fig. 6C- E**). Collectively, these results suggest that TNFα/TNFR1 signalling plays a major role in driving FLIP dependence in mutant Kras-driven lung cancers.

**Fig. 6:**
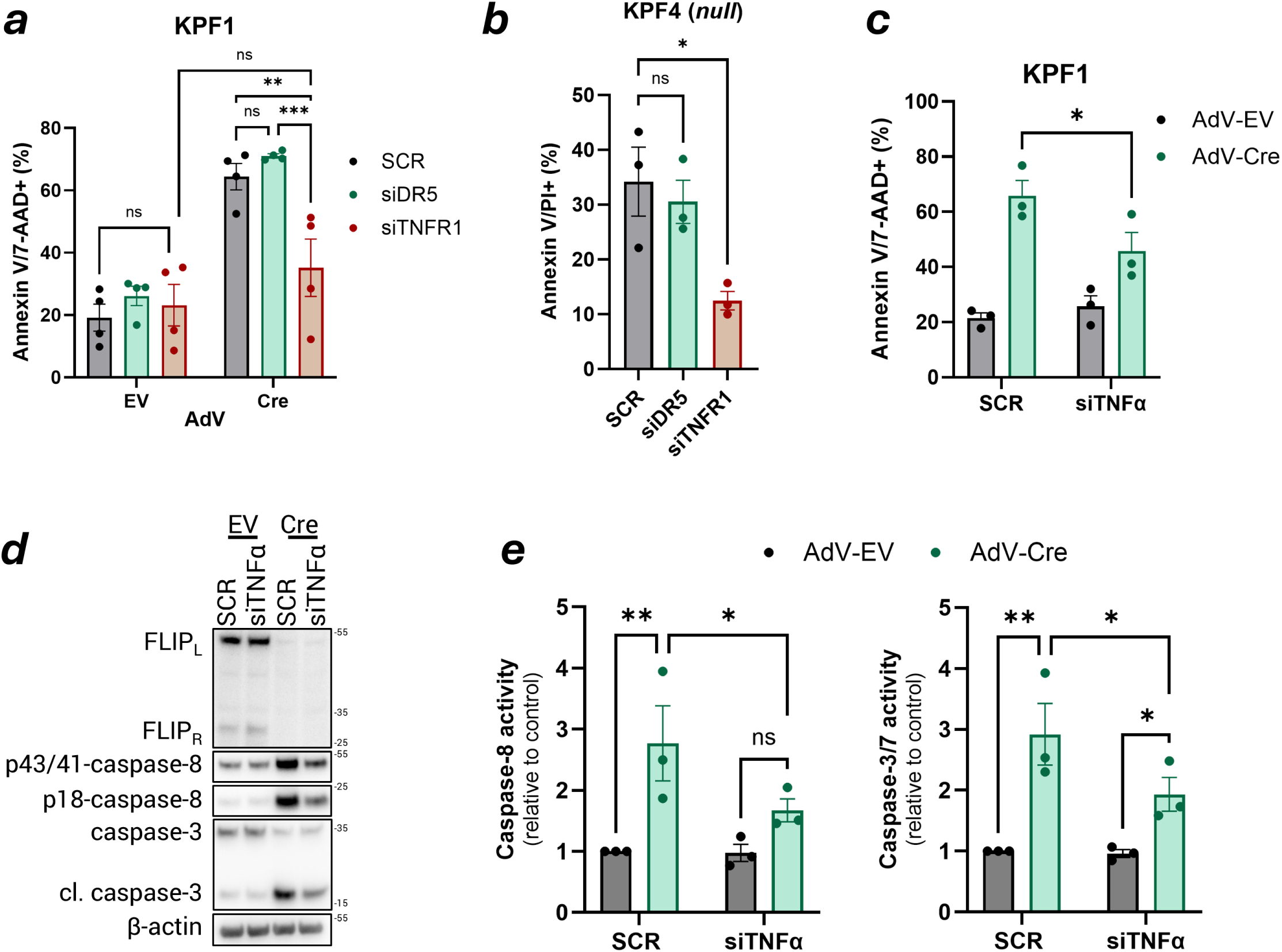
FLIP-deficiency drives TNFα /TNFR1-mediated apoptosis in *Kras* lung cancer. (**A**) Annexin V/7-AAD analysis in KPF1 cells following simultaneous transduction with Cre recombinase (AdV-Cre) or empty vector (AdV-EV) and transfection with control (SCR), DR5 or TNFR siRNA for 48 hours (n=4) Statistical tests were performed using two-way ANOVA and adjusted for multiple comparisons with Bonferroni correction. (**B**) Annexin V/PI analysis in KPF3 and KPF4 cells transfected with control (SCR) DR5 or TNFR siRNA for 48 hours (n=3). Results were compared using one-way ANOVA and adjusted for multiple comparisons with Dunnetts correction. (**C**) Annexin V/7-AAD analysis in KPF1 cells following simultaneous transduction with AdV-Cre or AdV-EV and transfection with control (SCR) or TNFα siRNA for 48 hours (n=3). Statistical tests were performed using two-way ANOVA and adjusted for multiple comparisons with Bonferroni correction. (**D**) Western blot analysis of FLIP_L_, FLIP_R_, p43/p18-caspase-8, caspase-3 and β- actin in KPF1 cells following treatment as in (B). (**E**) The activity of caspase-8 and caspase-3/7was analysed in extracted protein lysates from cells treated as in (**D**) using CaspaseGlo® activity assays, (n=3). Statistical tests were performed using two-way ANOVA. Data are mean +/- SEM. (\**p <* 0.05, \*\**p <*0.01, ***p<0.001,**** p<0.0001, ns – non-significant).

### Mutant *Kras* upregulates FLIP transcription via ERK1/2

To determine whether mutant *KRAS directly* contributes to elevated FLIP expression in *KRAS* mutant disease, we assessed the impact of the KRAS G12D inhibitor MRTX1133 (KRASi) on FLIP expression. In both the KPF1 and KPF2 cell line models, KRASi treatment resulted in downregulation of FLIP mRNA expression (**Fig. 7A and Supplementary Fig. 5A**), indicating that mutant KRAS positively regulates FLIP expression at the transcriptional level. This was reflected in time- dependent decreases in expression of both FLIP(L) and FLIP(R) proteins, with levels reaching ∼50% of control by 24h (**Fig. 7A and Supplementary Fig. 5B**). FLIP downregulation was preceded by decreases in pERK1/2 and pS6 kinase (**Fig. 7B and Supplementary Fig. 5B**) and correlated with increased cleavage (**Fig. 7B and Supplementary Fig. 5B**) and activity (**Fig. 7C and Supplementary Fig. 5C**) of caspase-8. Furthermore, enhanced caspase-8 activity correlated with enhanced activity of the executioner caspases-3/7 (**Fig. 7D and Supplementary Fig. 5D**). q- PCR and Western blot analyses of KPF1 cells treated with ERK and S6 kinase inhibitors indicated that the effects of mutant KRAS on FLIP expression are predominantly mediated through ERK1/2 (**Fig. 7E/F**). Notably, KRASi-induced apoptosis in KPF cells was found to be caspase-8-dependent (**Fig. 7G/H and Supplementary Fig. 5E**), and KRASi treatment failed to enhance apoptosis in the FLIP KO models (**Supplementary Fig. 5F**), collectively indicating that FLIP down- regulation and caspase-8 activation are key mechanisms by which oncogenic KRAS inhibition induces apoptosis. Consistent with its effects on FLIP expression, KRASi treatment resulted in enhanced sensitivity to TNFα- and TRAIL-induced apoptosis (**Fig. 7I**).

**Fig. 7:**
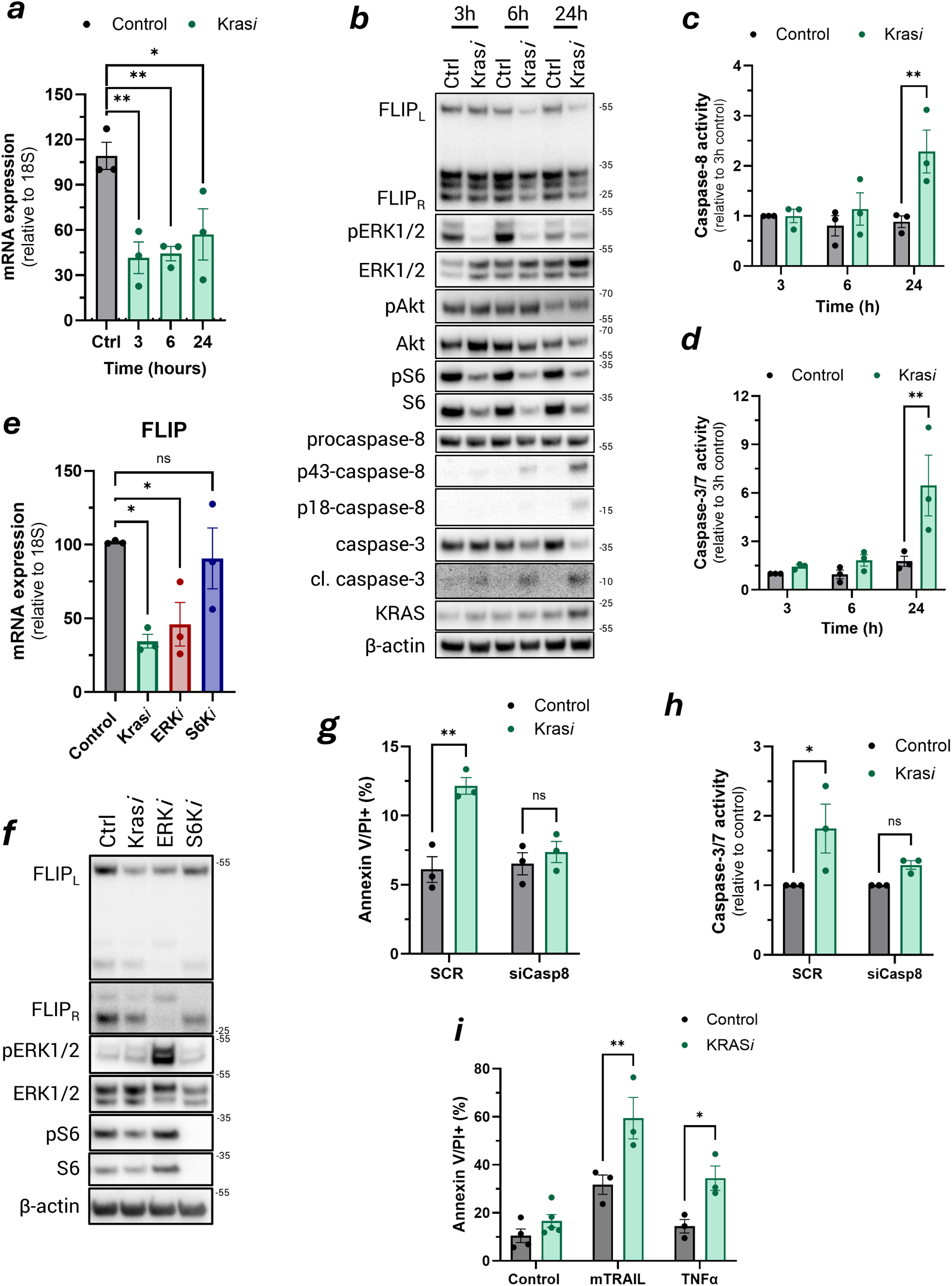
Mutant *Kras* upregulates FLIP transcription via ERK1/2. (**A**) Expression of FLIP mRNA (relative to 18S) in KPF1 cells following treatment with KRAS inhibitor MRTX1133 (300nM) or DMSO control for 3, 6 and 24 hours. Significance was tested using one-way ANOVA and adjusted for multiple comparisons with Dunnetts correction. (**B**) Western blot analysis of FLIP_L_, FLIP_R_, pERK1/2, ERK1/2, pAkt, Akt, pS6, S6, p43/p18-caspase-8, caspase-3, KRAS and β-actin in KPF1 cells following treatment with 300nM MRTX113 for 3, 6 and 24 hours. The activity of caspase-8 (**C**) and caspase-3/7 (**D**) was analysed in extracted protein lysates from cells treated as in (B) using CaspaseGlo® activity assays (n=3). (**E**) Expression of FLIP mRNA (relative to 18S) in KPF2 cells following treatment as in (**E**) (n=3). Significance was tested using one-way ANOVA and adjusted for multiple comparisons with Dunnetts correction. (\**p <* 0.05, ns – non-significant). (**F)** Western blot analysis of FLIP_L_, FLIP_R_, pERK1/2, ERK1/2, pAkt, Akt, pS6, S6, p43/p18-caspase-8, caspase-3, KRAS and β-actin in KPF2 cells following treatment with 300nM MRTX113, 1µM ERK inhibitor (LY3214996), S6K inhibitor (PF4708671) or DMSO (ctrl) for 24 hours. (**G**) Annexin V/PI analysis in KPF1 cells transfected with control (SCR) and Casp8 siRNA for 24 hours prior to treatment with 300nM KRAS*i* (MRTX1133) for a further 24 hours. Statistical tests were performed using two-way ANOVA and adjusted for multiple comparisons with Bonferroni correction. (**H**) Caspase-3/7 activity was analysed in extracted protein lysates from cells treated as in (**E**) using CaspaseGlo® activity assays, (n=3). (**I**) Annexin V/PI analysis of KPF1 cells pretreated with KRAS*i* MRTX1133 (300nM) for 24 hours prior to the addition of mTRAIL (25ng/mL) or TNFα (40ng/mL) for a further 24 hours. Statistical tests were performed using two-way ANOVA and adjusted for multiple comparisons with Bonferroni correction. Data are mean +/- SEM.

## DISCUSSION

*KRAS* is an oncogenic driver mutation found in ∼30% of NSCLC, as well as a significant proportion of other solid tumours, including CRC and pancreatic(3, 46, 47). Although mutant KRAS inhibitors have entered the clinic, both intrinsic and acquired resistance are substantial obstacles to the success of these agents(48). A deeper understanding of KRAS dependency and the mechanisms underlying response and resistance to these inhibitors is crucial for effective patient stratification and clinical benefit. Furthermore, identifying dependencies driven by oncogenic *KRAS* signalling is important so that the intrinsic vulnerabilities of these cancers are better understood and therapeutically exploited.(3, 46, 47).

We have previously reported that FLIP is frequently overexpressed in non-small cell lung cancer and that tumours with high FLIP expression have poor prognosis(37). These established tumour cell line models are dependent on FLIP expression, as its depletion promotes apoptosis independent of death ligand signalling(38). Here, utilising GEMMs of *Kras*-driven lung adenocarcinoma (*Kras^G12D/+^*/*Trp53^null^;* “KP”) in which the gene encoding FLIP, *Cflar*, could be conditionally knocked out, we show for the first time that the presence of FLIP is also essential for mutant Kras-driven lung tumourigenesis. Any tumours that did form after a prolonged latency period had (as expected) activated Kras^G12D^ and loss of p53, but had retained *Cflar*/FLIP, confirming that FLIP is a *sine qua non* for the growth of these tumours. Analyses of DepMap datasets demonstrated that FLIP mRNA and FLIP dependency are significantly higher in mutant *KRAS* lung cancers relative to wild-type, confirming that oncogenic KRAS-driven dependency on FLIP also occurs in human lung cancers. Indeed, these correlations held up across all solid tumour cell lines, with KRAS mutant models expressing higher FLIP and exhibiting higher FLIP-dependency, suggesting that these findings are likely relevant to other KRAS-driven cancers.

We explored the mechanisms behind FLIP-dependence in *Kras* mutant cancer by establishing cell lines (isolated from KPF animals) from “*Cflar* recombination escapee” tumours, *i.e.* tumours that grew by retaining the FLIP allele. We subsequently triggered *Cflar*/FLIP recombination *ex vivo* in these models. FLIP deletion resulted in caspase-8 and caspase-3/7 activation and cell death in KP lung cancer cells, confirming that FLIP is required to suppress cell death in *Kras* mutant lung cancer. We further demonstrated that this death is specifically controlled by caspase-8 activity. Interestingly, we found that caspase-8 mRNA expression is also elevated in human mutant *KRAS* lung cancer cell lines and positively correlates with FLIP dependence. Moreover, we show that KRAS inhibitor treatment downregulates FLIP expression at the mRNA and protein level and induces caspase-8-dependent apoptosis *in vitro*. Thus, the activation of oncogenic KRAS signalling drives a fundamental requirement for increased FLIP expression to inhibit caspase-8-mediated apoptosis. Oncogenic KRAS has previously been linked to control of apoptotic signalling through modulation of the intrinsic, mitochondrial-mediated apoptotic pathway by regulating expression of factors such as BCL2, BCLXL(49) and even through direct interaction with BAX and BAK (50); however, to our knowledge, this is the first time that oncogenic KRAS has been demonstrated to directly impact extrinsic apoptotic signalling.

Interestingly, despite elevated caspase activity and basal cell death rates, FLIP null KPF lung cancer cells retained colony-forming potential that was similar to that of FLIP WT KP cells. However, FLIP null KPF cells failed to establish lung tumours *in vivo* when transplanted into immune-competent mice, despite their ability to form colonies *in vitro*. These data confirm that KRAS mutant tumour cells remain dependent on FLIP beyond the tumour initiation stage, as these fully transformed FLIP null lung adenocarcinoma cell lines still cannot establish *in vivo*, suggesting that FLIP remains essential for *KRAS* mutant tumour growth in both early and late stages of lung cancer, influenced by both cell intrinsic and extrinsic factors. Since FLIP null KPF lung cancer cells were found to be hyper-sensitive *in vitro* to TRAIL, we assumed that TRAIL-expressing natural killer cells and CD8+ T lymphocytes would be responsible for the rejection of FLIP null KPF lung cells; however, these cells also failed to establish lung tumours in highly immunocompromised mice (C57BL/6N-*Rag2^Tm1^-IL2rg^Tm1^*/Rj) that lack these anti-tumour immune effector cells. Boumelha *et al.* reported similar findings when conducting *in vivo* CRISPR screening using a similar KRAS^G12D^ p53 null lung cancer model: they found that FLIP-targeting sgRNAs were equally depleted in immunocompetent and immunodeficient (*Rag2*^−*/*−^;*Il2rg*^−*/*−^) mice(51). They concluded that the *in vivo* requirement for FLIP must not be dependent on antitumour immunity; however, it is important to note that these immunocompromised mice retain functional macrophages capable of TNFα secretion(52). Moreover, KRAS mutations promote the recruitment of TNFα-secreting, proinflammatory macrophages to tumours(53). Furthermore, we found that Kras MT FLIP null lung cancer cells are highly sensitive to recombinant TNFα *in vitro* and that the TNFα/TNFR pathway is the critical mediator of cell death in these cells. In addition, KRAS inhibition enhanced exogenous TNFα-induced apoptosis *in vitro*. Together, these data suggest that mutant *Kras* cells that have lost FLIP are hypersensitive to TNFα and that the residual macrophage functionality in *Rag2*^−*/*−^;*Il2rg*^−*/*−^ mice may provide a source of TNFα *in vivo* that induces apoptosis and prevents lung colonisation of FLIP-null *Kras* mutant cells.

Interestingly, compared to their FLIP-proficient counterparts, FLIP-null lung cancer cells are also more sensitive to cisplatin, but not to agents directly targeting the intrinsic apoptotic pathway (BCL-2 (ABT199, ABT263), BCL-xL (ABT263) and MCL-1 (AZD5991)). These data indicate that cisplatin promotes death in a FLIP- dependent manner and suggest that FLIP expression might also be a marker of platinum sensitivity in *KRAS* mutant cancers. Indeed, our previous research has demonstrated that siRNA-mediated FLIP depletion synergises with cisplatin treatment(37, 38). KRAS inhibitors are often deployed in combination with chemotherapy, typically platinum-based doublet treatment, in advanced lung tumours. The downregulation of FLIP expression by KRASi has the potential to synergise with platinum-driven apoptosis, enhancing clinical benefit of KRASi even in earlier-stage disease. Thus, FLIP expression could also be used clinically to guide combination chemotherapy choices.

Using specific inhibitors of the branches of oncogenic KRAS signalling, we identified the ERK1/2 pathway to be primarily responsible for FLIP downregulation. This suggests that downregulation of FLIP in response to ERK inhibition will also be an important mechanism-of-action of MEK/ERK targeted therapeutics. This is increasingly relevant for KRASi acquired resistance, which often involves reactivation of MAPK signalling pathways via multiple mechanisms(54). Our results suggest that reactivation of ERK signalling in the KRASi-resistant setting will likely reinstate expression and dependency on FLIP, the expression of which could even be an early marker for acquired resistance.

Collectively, these novel results demonstrate that *KRAS* mutation confers sustained dependence on FLIP for tumour development and progression. Not only does this identify FLIP as an excellent therapeutic target in these cancers, but it also identifies a hitherto overlooked mechanism by which KRAS inhibitors exert their anti-tumour activity and highlight FLIP’s potential utility in stratifying patients to enhance the clinical efficacy of these therapeutics.

## Acknowledgements

This work was supported by grants from the MRC (MR/S021205/1), BBSRC (BB/T002824/1) and Wellcome (110371/Z/15/Z). Emma Kerr was supported by a CRUK Career Development Award (C61288/A26045). We thank Kirsty McLaughlin for technical support and flow cytometry expertise and Prof Owen Sansom for the Kras mouse model. We thank the Biological Services Unit at QUB for technical support with in vivo experiments.

## Completing interests

DBL, TH and CH are in receipt of funding from Ipsen Global for development of inhibitors of FLIP, although these were not used in this study.

## Materials and Methods

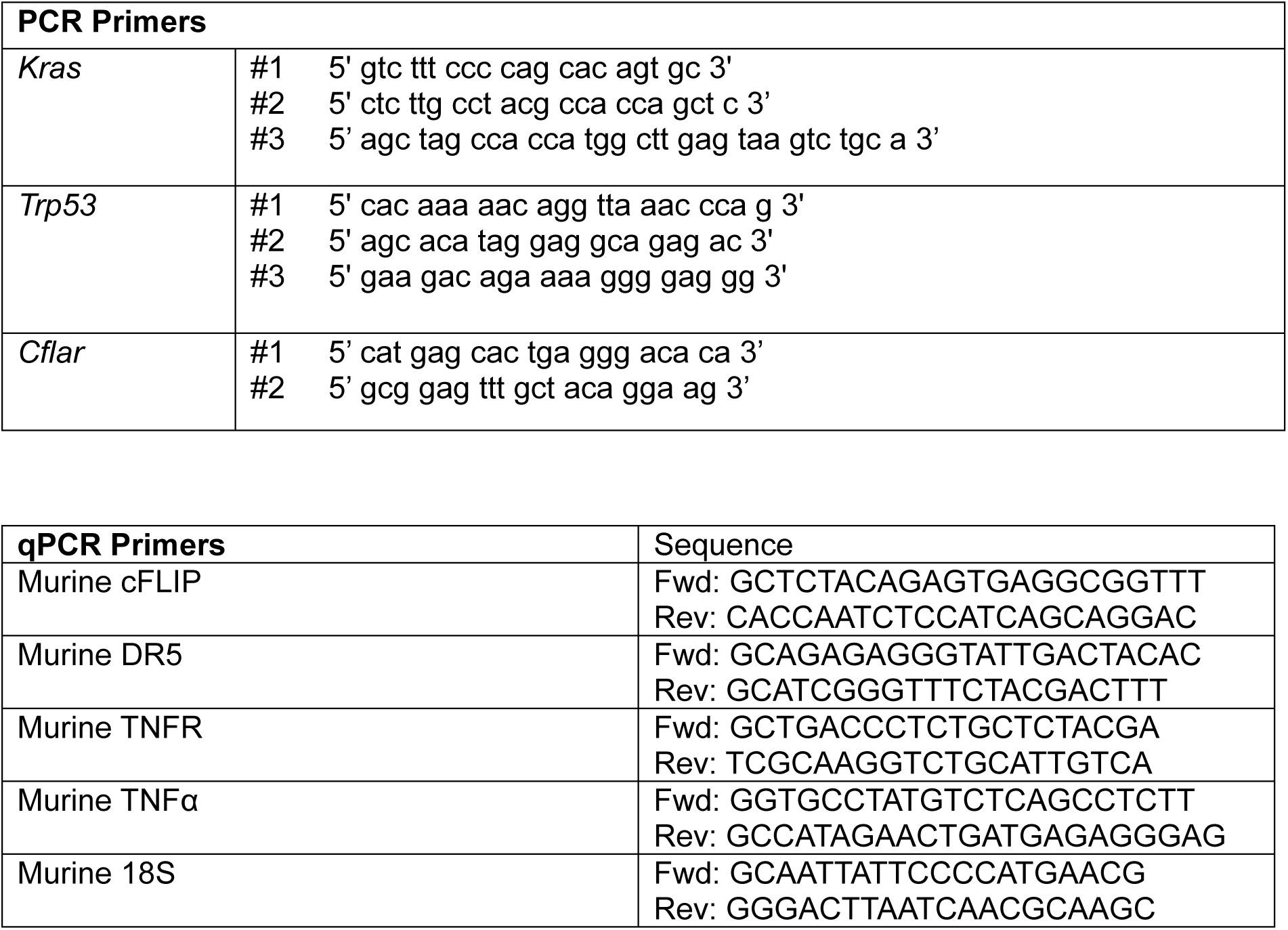

## Supplementary Figures

**Supplementary Fig. 1.**
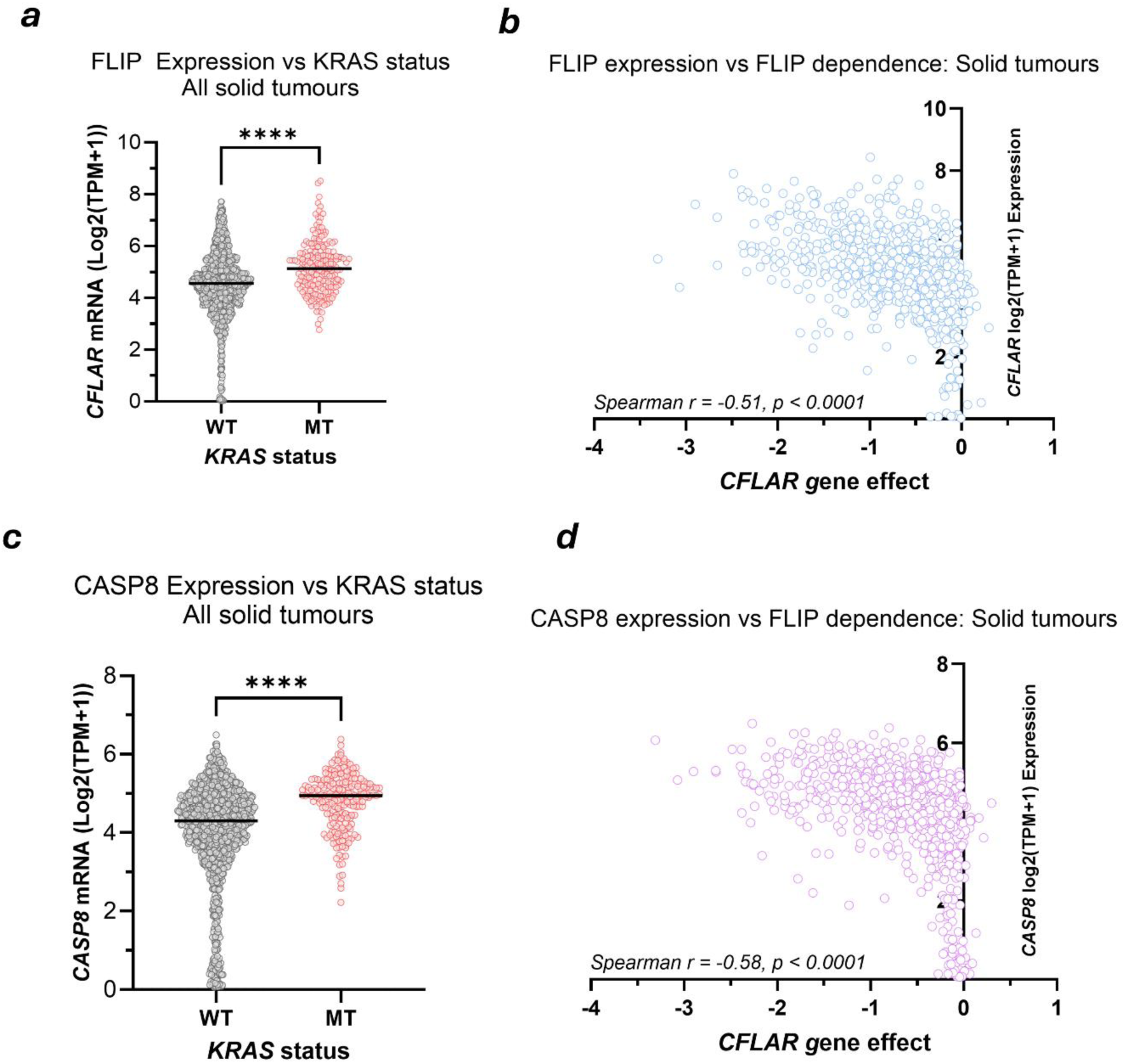
(**A**) Expression of FLIP/*CFLAR* mRNA in *KRAS* WT and MT solid cancer cell lines in the DepMap panel of solid tumour cell lines. (**B**) Correlation of FLIP/*CFLAR* gene expression with FLIP/*CFLAR* gene dependency in solid cancer cell lines in the DepMap panel. (**C**) Expression of Caspase-8/*CASP8* mRNA in *KRAS* WT and MT solid cancer cell lines in the DepMap panel. (**D**) Correlation of Caspase-8/*CASP8* gene expression with FLIP/*CFLAR* gene dependency in solid cancer cell lines in the DepMap panel.

**Supplementary Fig. 2.**
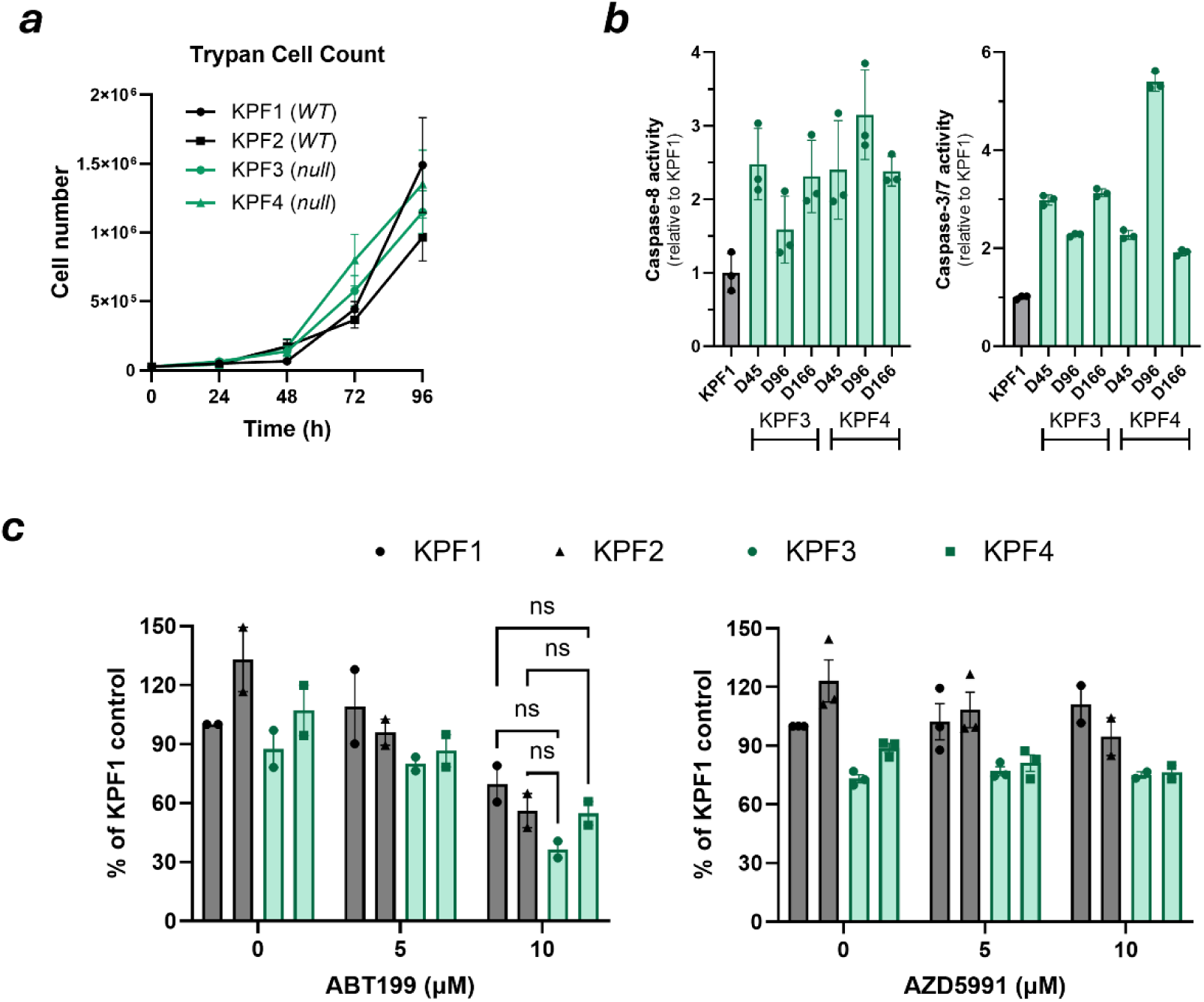
(**A**), Growth curves of KPF1-4 cells over time. Trypan blue stain was used to count the number of viable cells at 24-, 48-, 72- and 96-hour timepoints in three independent experiments. (**B**) Baseline caspase-8 and caspase-3/7 activity in KPF3 and KPF4 cells at day 45, 96 and 166 after the removal of AdV-Cre. KPF1 was used as a control. (**C**) Cell viability assays in KPF1-4 cell lines following treatment with increasing doses (1.25, 2.5, 5 and 10µM) of BH3 mimetics ABT199 and AZD5991. Data were normalized to KPF1 control, (n>=2). Statistical tests were performed using two-way ANOVA and adjusted for multiple comparisons with Bonferroni correction. (ns – non-significant). Data are mean +/- SEM.

**Supplementary Fig. 3.**
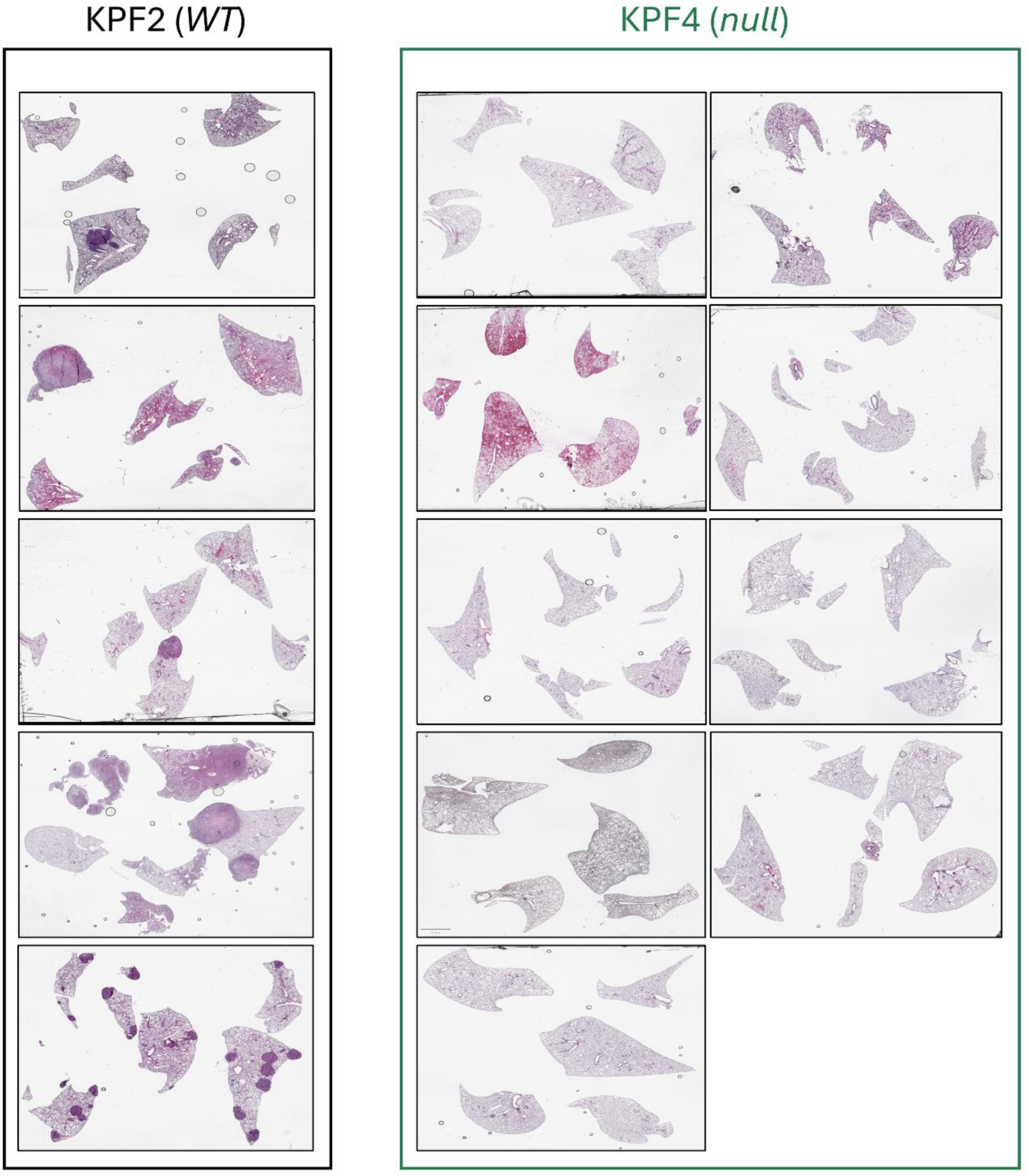
H&E of lung sections from C57Bl/6J mice after 13 weeks post-inoculation with KPF2 (*WT*) and KPF4 (*null*) cells (Fig. 4 **(F)**) and used for quantification of histologic tumour burden (tumour area/total lung area) (Fig. 4**(H)**).

**Supplementary Fig. 4.**
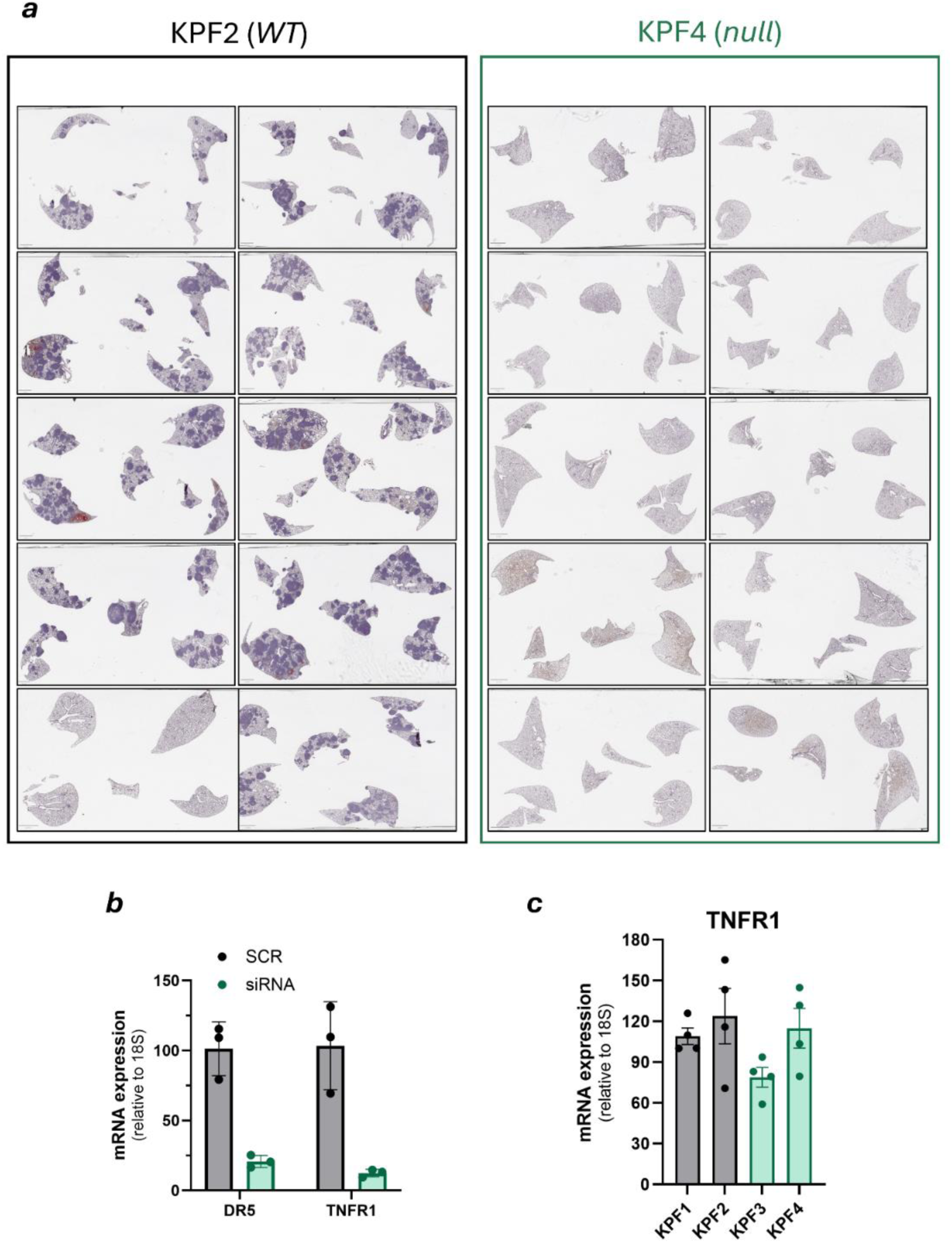
(**A**) H&E of lung sections from C57BL/6N-Rag2Tm1-IL2rgTm1/Rj mice after experimental endpoint following inoculation with KPF2 (*WT*) and KPF4 (*null*) cells (Fig. 5 **(A)**) and used for quantification of histologic tumour burden (tumour area/total lung area) (Fig. 5 **(C)**). (**B**) Expression of DR5 and TNFR mRNA in KPF1 cells following treatment as in Fig. 5 **(F)**. (**C**) Expression of TNFR mRNA (relative to 18S) in KPF1-4 cells, (n=4).

**Supplementary Fig. 5.**
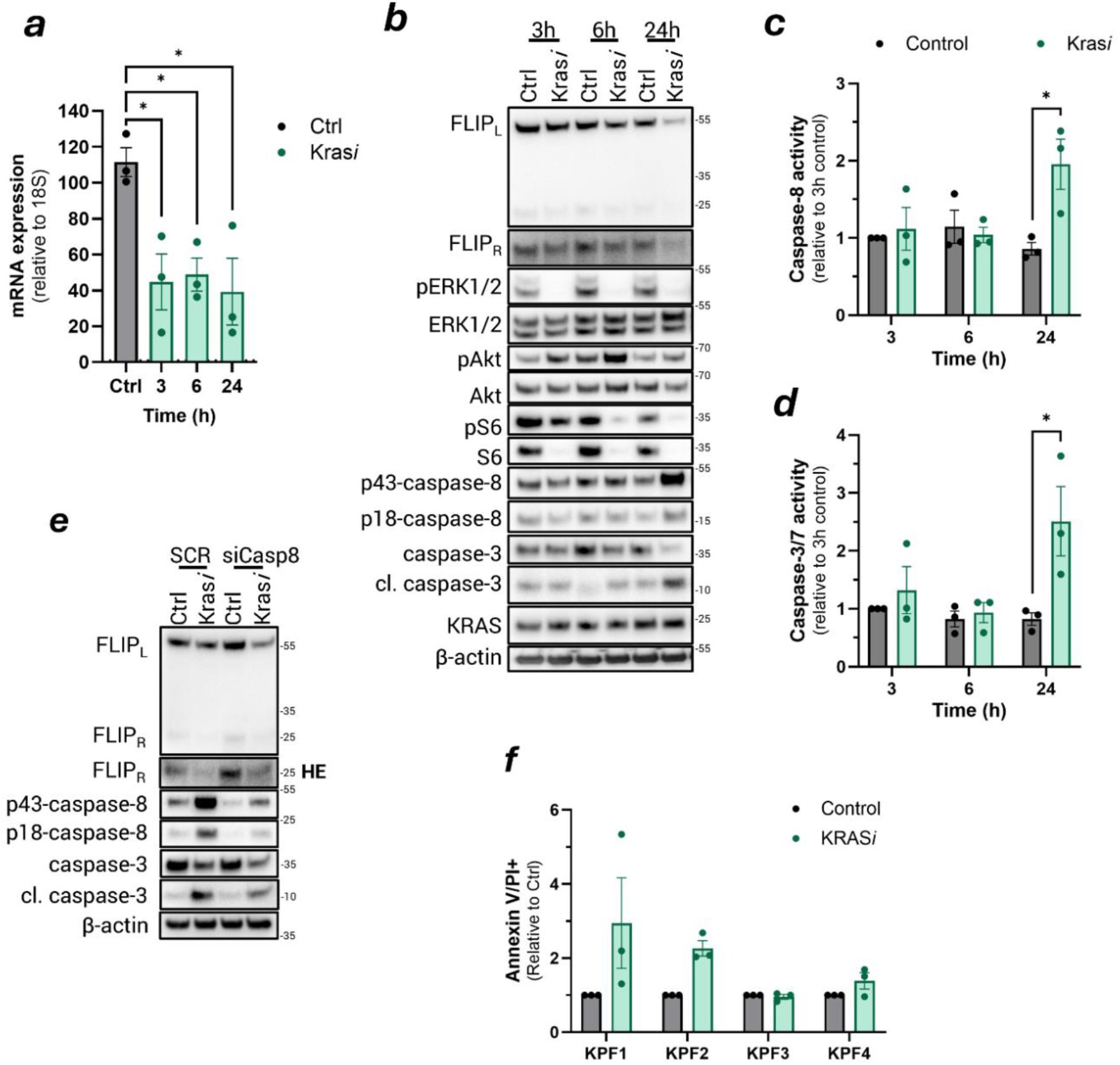
(**A**) Expression of FLIP mRNA (relative to 18S) in KPF2 cells following treatment with KRAS*i* MRTX1133 (300nM) or DMSO control for 3, 6 and 24 hours (n=3). Significance was tested using one-way ANOVA and adjusted for multiple comparisons with Dunnetts correction. (**B**) Western blot analysis of FLIP_L_, FLIP_R_, pERK1/2, ERK1/2, pAkt, Akt, pS6, S6, p43/p18-caspase-8, caspase-3, KRAS and β-actin in KPF2 cells following treatment with 300nM of KRAS*i* MRTX113 for 3, 6 and 24 hours. The activity of caspase-8 (**C**) and caspase-3/7 (**D**) was analysed in extracted protein lysates from treated as in (**B**) using CaspaseGlo® activity assays (n=3). Results were compared using Students *t*-test. (**E**) Western blot analysis of FLIP_L_, FLIP_R_, p43/p18-caspase-8, caspase-3 and β-actin in KPF1 cells following treatment as in Fig. 7(G). (**F**) Annexin V/PI analysis in *Cflar WT* (KPF1 and KPF2) and *null* (KPF3 and KPF4) cells following treatment with 300nM KRAS*i* (MRTX1133) for 24 hours (n=3). Data is shown as fold change relative to control for each cell line. Data are mean +/- SEM.

